# Swirling Instability of the Microtubule Cytoskeleton

**DOI:** 10.1101/2020.08.27.268318

**Authors:** David B. Stein, Gabriele De Canio, Eric Lauga, Michael J. Shelley, Raymond E. Goldstein

## Abstract

In the cellular phenomena of cytoplasmic streaming, molecular motors carrying cargo along a network of microtubules entrain the surrounding fluid. The piconewton forces produced by individual motors are sufficient to deform long microtubules, as are the collective fluid flows generated by many moving motors. Studies of streaming during oocyte development in the fruit fly *D. melanogaster* have shown a transition from a spatially-disordered cytoskeleton, supporting flows with only short-ranged correlations, to an ordered state with a cell-spanning vortical flow. To test the hypothesis that this transition is driven by fluid-structure interactions we study a discrete-filament model and a coarse-grained continuum theory for motors moving on a deformable cytoskeleton, both of which are shown to exhibit a *swirling instability* to spontaneous large-scale rotational motion, as observed.

---

A striking example of fluid-structure interactions within cells [1] occurs in oocytes of the fruit fly *Drosophila melanogaster* [2]. These develop over a week from a single cell through repeated rounds of cell division, differentiation and growth, ultimately reaching hundreds of microns across. This pathway has historically been divided into 14 stages, and it is in stages 9 — 11, at days 6.5 — 7 [3], that fluid motion is most noticeable. In stage 9 (Fig. 1), microtubules (MTs) reach inward from the oocyte periphery, forming a dense assembly along which molecular motors (kinesins) move at tens of nm/sec, carrying messenger RNAs and other nanometric particles. This motion entrains the surrounding fluid, producing cytoplasmic streaming [4, 5] that can be visualized several ways: in brightfield by the motion of endogenous particles [6–8], via their autofluorescence [9, 10], and through a combination of particle image velocimetry and fluorescently labelled microtubules [11–13]. Previous work [7, 11] revealed that these flows initially take the form of transient, recurring vortices and jets whose correlation length is a fraction of the cell scale, with no long-range order. But by stage 11, a dramatic reconfiguration of the cytoskeleton occurs, coincident with the appearance of a vortex spanning the entire cell [6, 7, 10, 14].

**FIG. 1.**
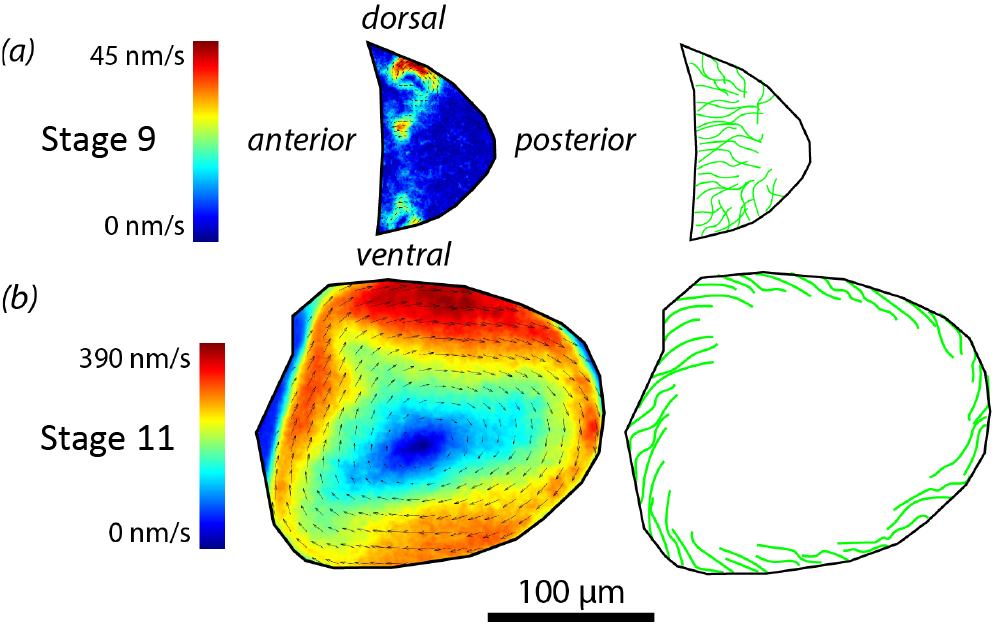
Cytoplasmic streaming flows in the *Drosophila* oocyte. (a) Experimental flow field [28] and schematic of the disordered swirling flows and microtubule organization in early stages of development. (b) Later flows organize into a single vortex as MTs lie parallel to the cell periphery.

Kinesin motors move from the *minus* ends of microtubules (attached to the oocyte periphery) to the *plus* ends (free in the interior). Transport of cargo through the network depends on motor-microtubule binding details [15, 16] and on the mesh architecture [17, 18]. As a motor pulls cargo toward the plus end the filament experiences a localized minus-end-directed compressive force, as in Euler buckling. For a filament of length *L* and bending modulus *A* [19], the buckling force is ~ *πA/L*^2^ ~ 60pN/*L*^2^, where L is measured in microns. Thus, a kinesin’s force of several pN [20] can buckle MTs 10 — 40 μm long.

The coupled filament-motor problem is richer than Euler buckling because a motor exerts a “follower force” [21] that is aligned with the filament. This feature breaks the variational structure of the problem and allows a filament pinned at its minus end to oscillate even at zero Reynolds number [22–24]. By exerting a force on the fluid a motor induces long-range flows which, if compressive, can further deform filaments [25, 26].

It has been hypothesized [10, 14] that the transition from disordered streaming flows to a single vortex in stage 11 is a consequence of the kinds of fluid-structure interactions described above, facilitated by a decrease in cytoplasmic viscosity that accompanies the disappearance of a coexisting network of the biopolymer f-actin. Here, through a combination of direct computations on the coupled filament-flow problem [23] and studies of a recent continuum theory for dense filament suspensions [27], we confirm this hypothesis by showing the existence of a novel *swirling instability* of the cytoskeleton.

The swirling instability can be understood in a simplified model of the oocyte: a rigid sphere of radius *R* containing a fluid of viscosity *μ*, with *N* elastic filaments reaching inwards from clamped attachment points equally spaced around the equator. A slice in the filament plane (Fig. 2(a)) appears like a confocal slice of the oocyte (Fig. 1). The filaments have a radius *r*, a constant length *L*, bending modulus *A* and a uniform line density *f* of follower forces (Fig. 2(b)). Although free microtubules have a complex dynamics of growth and decay, recent evidence [29] for ‘superstable’ cortically-bound microtubules in stages displaying unidirectional streaming justifies the constant-length approximation.

**FIG. 2.**
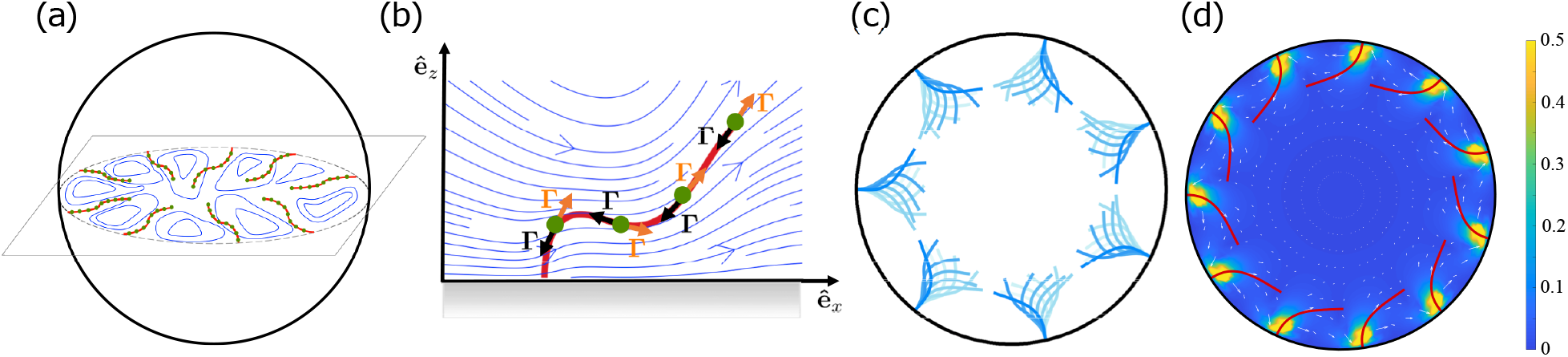
Discrete filament computations. (a) *N* equally spaced elastic filaments clamped at their attachment points, reach inward from a no-slip spherical shell. Each has a continuous distribution of tangential point forces (red) that (b) exert a force **Γ** on the fluid and an equal and opposite compressive force on the filament. Synchronous oscillations (*N* = 7, *σ* = 1700), (d) steady, bent configuration (*N* = 9, *σ* = 500) and swirling flow field.

Microtubules are the quintessential slender bodies [30] of biophysics, with aspect ratios *ε* = *r/L* of 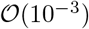. As their self-interactions are weak, we use local slender-body theory [31, 32] to obtain the dynamics. In an arclength parameterization *s*, the *j*^th^ filament **r**^*j*^(*s, t*) evolves as

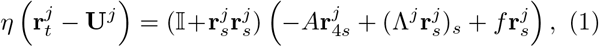

where 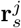 is the unit tangent, *η* = 8*πμ/c*, with *c* = |ln(*eε*^2^)|, and the Lagrange multiplier Λ^*j*^ enforcing inextensibility obeys a second-order PDE [33]. In the background flow **U**^*j*^ = **u**^*j*^ + **u**^*i→j*^ + **v**^*i→j*^, **u**^*j*^ is that produced by the motors on *j*, **u**^*i→j*^ is due to the motors on *i ≠ j*, and **v**^*i→j*^ is due to motion of filaments *i ≠ j*. For example, the induced flow due to the *j*th fiber is 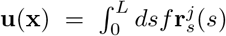 (see Supplemental Material [34, 35]), with **G** the Greens function appropriate to the interior of a no-slip sphere [36]. Filament clamping at the sphere implies that **r**^*j*^ (0, *t*) remains fixed and that 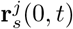 is the inward sphere normal at the attachment point. The free end is torque- and force-free: 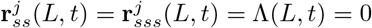.

A single fiber clamped at a flat wall displays a supercritical Hopf bifurcation which, expressed in terms of the dimensionless motor force *σ* ≡ *fL*^3^/*A*, occurs at *σ*\* ≃ 124.2, beyond which the filament exhibits steady oscillations whose amplitude grows as 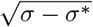 [23]. When several filaments interact within the sphere (2c) they also oscillate, but with their motions synchronized in phase, very much like eukaryotic flagella [37]. The dynamical model (1) contains two ingredients often found necessary for such synchronization [38]: hydrodynamic interactions and the ability of a filament to change shape and thereby adjust its phase in response to those flows.

As the filament density and motor strength are increased we find the swirling instability: a transition to a *steady* configuration of bent filaments whose distal parts are oriented almost parallel to the wall (Fig. 2(d)). The bent configuration is maintained by azimuthal flows, induced by the motors, that generate drag along the distal part of the filaments, and thus a torque opposing the bending torques near the filament base. As with any such spontaneous breaking of symmetry, both left- and right-hand configurations are possible; the choice between the two is dictated by initial conditions. This transition is reminiscent of the self-organized rotation of cytoplasmic droplets extracted from plants [39] and the spiral vortex state of confined bacterial suspensions [40], both modeled as suspensions of stresslets [41-43].

While direct computations on denser arrays of discrete filaments are possible [44], cortically bound oocyte microtubules are so tightly packed, with an inter-fiber spacing *δ ≪ L* [10-13], that a continuum approach is justified. The description we use [27], in which microtubules form an anisotropic porous medium, is based on the map **X** = **r**(***α***), where the Lagrangian coordinate **α** = (*α, s*) encodes the location *α* of the minus ends of the microtubules and arclength *s*. In a system of units made dimensionless by the length *L* and elastic relaxation time *ηL*^4^/*A*, we obtain a continuum version of (1),

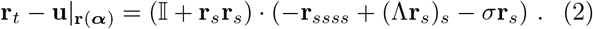

The fluid velocity **u** arises from the force distribution along the filaments and is evaluated at the Eulerian position **x** according to an inhomogeneous Stokes equation,

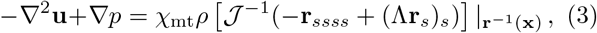

subject to the incompressibility constraint ∇ · **u** = 0. The indicator function *χ*_mt_ is supported where the MT array is present (Fig. 3a). Here, *ρ* = 8*πρ*_0_*L*^2^/*c* is the rescaled areal number density of microtubules, expressible as *ρ = ϕ*(*L/δ*)^2^, where the constant *ϕ* depends only on the MT slenderness and packing geometry at the wall; *ϕ* ≈ 4 when *c* ≈ 10 and the MTs are hexagonally packed. The quantity 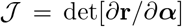 measures the change in microtubule density due to deformations of the array; 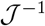 increases as fibers move closer together.

**FIG. 3.**
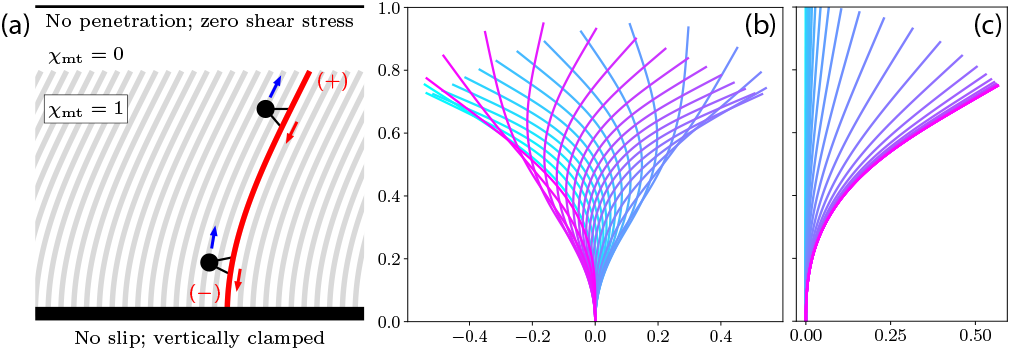
Continuum model in planar geometry. (a) Bi-infinite array of MTs, whose minus ends are clamped vertically at a no-slip boundary. Results of full computations at (b) *ρ* = 4.65, *σ* = 70 and (c) *σ* = 39. Colors denote time, from cyan (early) to pink (late).

The simplest geometry is an infinite planar array of MTs, as shown in Fig. 3(a). As in the discrete model, the MTs are normally clamped to a no-slip wall and are force- and torque-free at their plus ends. At a distance *H* above the wall, no-penetration and zero-tangential stress conditions are imposed on the fluid. For dynamics homogeneous along *x*, the fluid flow is unidirectional and constant above the MTs, so *H* plays no role. Nonlinear computations [45] reveal both oscillatory dynamics and the emergence of steady streaming. Fig. 3(b) shows the dynamics when *ρ* = 4.65 and *σ* = 70: self-sustaining oscillations of the MT array are observed, similar to those in Fig. 2(c). Note that while Fig. 3(b) shows only a single filament, it represents the common dynamics of *all* of the collectively beating filaments in the array. When σ is decreased to ≈ 39, the MT array deforms and stabilizes into a *steady* bent state (Fig. 3(c)). This represents the continuum description of the swirling transition, with similar dynamics to those observed in the discrete results.

An equilibrium of the system occurs when filaments are aligned straight along *z*, with **u** = 0 and **Λ** = −*σ*(1 − *z*). For *σ* > 0, the motor-force is compressive and buckling may occur. A small transverse perturbation in fiber shape of the form 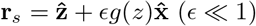 evolves as

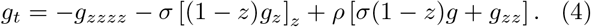

The first two terms are like those of an elastic filament under an aligned gravitational load, with an internal tension varying linearly from one end to the other [46, 47]. The third is the fiber forces filtered through the non-local Stokes operator, capturing hydrodynamic interactions within the fiber array (and hence the *ρ* prefactor). That this term is local is both fortunate and surprising, and follows from the simplicity of the Stokes flow in this case. The term *ρg_zz_* captures the additional resistance to bending from flow: if a MT is to bend, it must move the fluid around it, bending other MTs; the term *ρσ*(1 − *z*)*g* is destabilizing: if a MT is to remain straight, it must resist the fluid motions generated by MTs around it.

While the planar geometry reproduces all qualitative features of the streaming transition [34], to capture the key feature of confined hydrodynamic interactions in the oocyte we extend the analysis to a cylindrical domain, where the no-flow steady state is an array of straight MTs pointing inwards. Fig. 4 shows the results of a linear stability analysis for an experimentally relevant ratio of cylinder diameter to MT length of 10: 1.

**FIG. 4.**
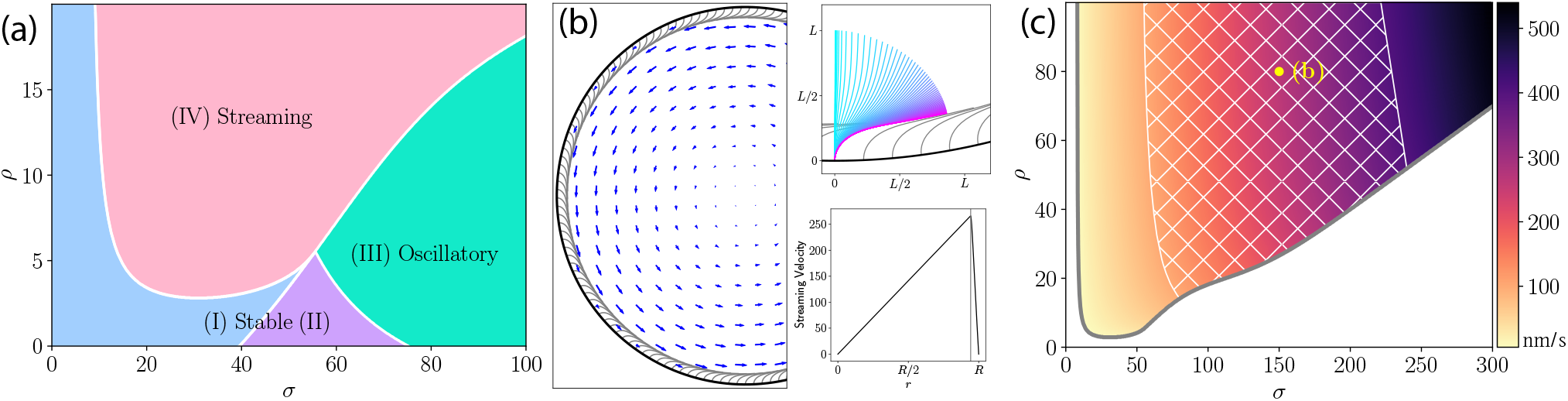
Continuum model in cylindrical geometry. (a) Results of linear stability analysis about the radially aligned state, with *R* = 5*L*. (b) Steady-state fiber deformations and velocity field for *σ* = 150 and *ρ* = 80. Density of visualized fibers corresponds to the physical density. Top inset shows deformed MTs and the dynamics of a representative one (see also supplemental video [34]). Bottom inset shows the azimuthal velocity field as a function of *r*. (c) Dimensional streaming velocities in parameter space; hatched region is consistent with *in vivo* estimates of 100 − 400nm/s. Yellow dot denotes simulation shown in (b).

For *ρ* ≪ 1, the continuum model behaves like isolated fibers with negligible collective fluid entrainment. For small *σ*, straight fiber arrays are stable (regions I & II, with region II having oscillatory decay to equilibrium), but with increasing *σ* there is a Hopf bifurcation to a state that nonlinear simulations show has oscillations (cf. Fig. 2(c)). For *ρ* > 2.8 (*δ* < 1.2L), a new region of instability (IV) appears, with real and positive eigenvalues; nonlinear simulations show this leads to collective MT bending and swirling flows.

Figure 4(b) shows a nonlinear simulation of the transition to streaming in region IV. The upper inset shows the development of the instability, with successive MTs bending over to form a dense canopy above their highly curved bases. At steady state, the concentrated motor forces within the canopy are azimuthally aligned, almost a *δ*-function a distance ~ *L*/4 above the wall, and drive the large-scale streaming flow. The ooplasmic flow beneath the MT canopy is nearly a linear shear flow, transitioning above to solid body rotation, the solution to Stokes flow forced at a cylindrical boundary.

We now estimate ranges of density and force that are consistent with observed streaming speeds *u* ≈ 100 − 400nm/s (Fig. 1 and [14, 29]). Taking *L* = 20*μ*m, *μ* =1 Pa s [11] and A = 20pN*μ*m^2^, we obtain a velocity scale *A/ηL*^3^ ≈ 1nm/s and a force-density scale *A/L*^3^ ≈ 2.5fN/*μ*m. Figure 4c shows the streaming speeds calculated from nonlinear simulations in region IV. Those with maximum speeds falling in the experimental range lie in the hatched area. Increasing *ρ* only marginally increases streaming speeds, and so to increase flow speed while remaining in region IV requires increasing both *ρ* and *σ*. The minimum value of *ρ* ≈ 20 that is consistent with observed streaming velocities corresponds to 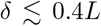, a more stringent constraint than that required for the streaming transition. The force densities consistent with streaming speeds are *f* ~ 0.1−0.6pN/*μ*m. Speeds on the higher end of the physical range approach the ≈ 700nm/s of kinesin-1 under negligible load [20], while cargo speeds on oocyte MTs are 200 − 500nm/s [14, 29, 48]. Assuming a linear force-velocity relation and a stall force of 6 pN [20] gives a single motor force of − 2 pN; approximately 1 − 6 kinesins are required per 20 *μ*m MT to generate these force densities.

It may be surprising that the streaming speed only weakly depends on *ρ*. A heuristic argument for the flow speeds views the cytoskeleton as a porous medium of permeability *k ~ δ*^2^, in which speed *u* ~ (*k/μ*)∇*p*, where the pressure gradient (force/volume) from motors is *f/δ*^2^, yielding *u ~ f/μ* ~ (*A/ηL*^3^)(8*π/c*)*σ*, independent of *ρ*. This relationship is surprisingly accurate [34].

When the density *ρ* is sufficiently high, the swirling instability first appears for force densities *σ* substantially smaller than those that induce buckling instabilities in a single filament. Thus this transition must be driven by the additional hydrodynamic destabilization that neighboring fibers impart (in the simplest geometry, given by the term *ρσ*(1 − *z*)*g* in Eq. 4). This observation motivates a simple heuristic argument for the instability, in which a filament is bent by the flow produced primarily by its upstream neighbor, whose distal half is nearly parallel to the wall. Seen from a distance, that bent portion acts on the fluid like a point force [49] **F** ~ (*fL*/2)*r_s_*(*L*) oriented along its distal tangent vector (Fig. 5), displaced a distance *h ~ L*/2 from the surface. Near a no-slip wall, the far-field flow along *x* due to a force 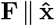 a distance *δ* upstream is simple shear [50, 51],

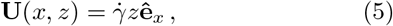

where 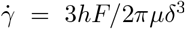. Self-consistency requires the magnitude of the force driving the shear be given by the projection of **F** along *x*, so 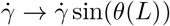.

**FIG. 5.**
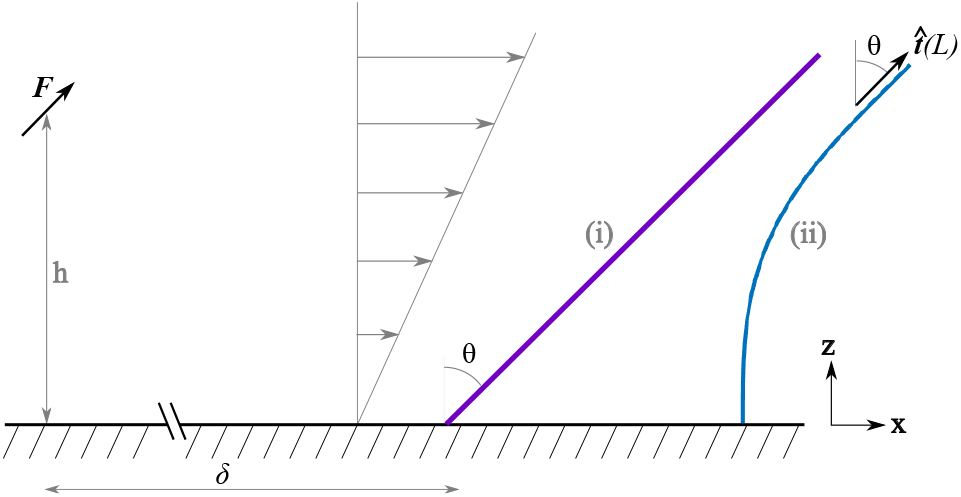
Self-consistent model. An upstream point force F parallel to the distal end of a filament produces shear ow that deects the filament. Two variants of the model: (i) rigid rod with a torsional spring at its base, (ii) a clamped elastic filament.

The very simplest model to illustrate the self-consistency condition is a rigid MT with a torsional spring at its base that provides a restoring torque −*kθ* (Fig. 5(i)). With *z*(*s*) = *s* cos *θ* and 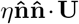 the local normal force on a segment, the local torque about the point *s* = 0 is 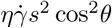 which, when integrated along the ament and balanced against the spring torque, yields the self-consistency condition

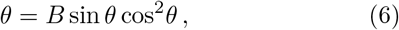

where 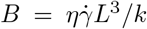. For *B* < 1 (slow flow or a stiff spring) *θ* = 0 is the only fixed point, while for 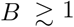 two mirror-image swirling solutions appear through a pitchfork bifurcation, *θ*_±_ ≃ (6(*B* − 1)/7)^1/2^.

To study the interplay between filament oscillations and swirling we use (5) in the filament dynamics (1), where the control parameter for the shear flow is [25, 26]

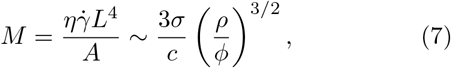

where the second relation uses the above estimates for *F* and *h*. Since a clamped elastic filament behaves like a torsional spring with a spring constant *k = A/L*, we see consistency with the parameter *B* defined above. A numerical self-consistent calculation confirms the existence of a swirling instability [34].

Through simplified discrete and continuum models we have demonstrated the existence of a novel swirling instability of arrays of elastic filaments, thus lending support to the hypothesis [14] that cytoplasmic streaming flows in *Drosophila* oocytes are tied to self-organization of the microtubule cytoskeleton. Future studies could shed light on the detailed mechanism involved in the untangling of the *Drosophila* oocyte cytoskeleton when it transitions to the vortical state, and the possibility of reproducing this transition *in vitro*. Lastly, this study highlights the role of active force dipoles in the self-organization of fluid-biopolymer systems [41–43].

We are indebted to Maik Drechsler and Isabel Palacios for sharing the data in Fig. 1 and to them, Daniel St Johnston and Vladimir Gelfand for discussions on *Drosophila* streaming. This work was supported in part by ERC Consolidator grant 682754 (EL), Wellcome Trust Investigator Award 207510/Z/17/Z, Established Career Fellowship EP/M017982/1 from the Engineering and Physical Sciences Research Council, and the Schlumberger Chair Fund (REG). MJS acknowledges the support of NSF Grant DMS-1620331.

## Reference

[1] D. Needleman and M.J. Shelley, The stormy fluid dynamics of the living cell, Physics Today 72, 32 (2019).

[2] A short primer on the subject is: R. Bastock and D. St. Johnston, *Drosophila* oogenesis, Curr. Biol. 18, R1082 (2008).

[3] L. He, X. Wang, and D.J. Montell, Shining light on *Drosophila* oogenesis: live imaging of egg development, Curr. Op. Gen. & Dev. 21, 612 (2011).

[4] The phenomenon of streaming was first discovered in plants, as reported by B. Corti, Osservazione Micro-scopische sulla Tremella e sulla Circulazione del Fluido in Una Planto Acquaguola (Appresso Giuseppe Rocchi, Lucca, Italy, 1774).

[5] R.E. Goldstein and J.-W. van de Meent, A Physical Perspective on Cytoplasmic Streaming, Interface Focus 5, 20150030 (2015).

[6] H. Gutzeit and R. Koppa, Time-lapse film analysis of cytoplasmic streaming during late oogenesis of *Drosophila*, J. Embryol. Exp. Morphol. 67, 101 (1982).

[7] W. Theurkauf, S. Smiley, M. Wong, and B. Alberts, Re-organization of the cytoskeleton during Drosophila oogenesis: implications for axis specification and intercellular transport Development 115, 923 (1992).

[8] W.E. Theurkauf, Premature microtubule-dependent cytoplasmic streaming in cappuccino and spire mutant oocytes, Science 265 2093 (1994).

[9] I.M. Palacios and D. St. Johnston, *Kinesin light chain-*independent function of the *Kinesin Heavy Chain* in cytoplasmic streaming and posterior localisation in the *Drosophila* oocyte, Development 129:54735485 (2002).

[10] L.R. Serbus, B.J. Cha, W.E. Theurkauf, W.M. Saxton, Dynein and the actin cytoskeleton control kinesin-driven cytoplasmic streaming in *Drosophila* oocytes, Development 132, 3743 (2005).

[11] S. Ganguly, L.S. Williams, I.M. Palacios, and R.E. Goldstein, Cytoplasmic Streaming in *Drosophila* Oocytes Varies with Kinesin Activity and Correlates With the Microtubule Cytoskeleton Architecture, Proc. Natl. Acad. Sci. USA 109, 15109 (2012).

[12] M. Drechsler, F. Giavazzi, R. Cerbino, and I.M. Palacios, Active diffusion and advection in *Drosophila* oocytes results from the interplay of actin and microtubules, Nat. Comm. 8, 1520 (2017).

[13] M. Drechsler, L.F. Lang, L. Al-Khatib, H. Dirks, M. Burger, C.-B. Schonlieb and I.M. Palacios, Optical flow analysis reveals that Kinesin-mediated advection impacts the orientation of microtubules in the *Drosophila* oocyte, Mol. Biol. Cell 31, 1246 (2020).

[14] C.E. Monteith, M.E. Brunner, I. Djagaeva, A.M. Bielecki, J.M. Deutsch, and W.M. Saxton, A Mechanism for Cytoplasmic Streaming: Kinesin-Driven Alignment of Microtubules and Fast Fluid Flows, Biophys. J. 110, 2053 (2016).

[15] P. Khuc Trong, J. Guck, and R.E. Goldstein, Coupling of Active Motion and Advection Shapes Intracellular Cargo Transport, Phys. Rev. Lett. 109, 028104 (2012).

[16] L.S. Williams, S. Ganguly, P. Loiseau, B.F. Ng and I.M. Palacios, The auto-inhibitory domain and the ATP-independent microtubule-binding region of Kinesin Heavy Chain are major functional domains for transport in the *Drosophila* germline, Development 141, 176 (2014).

[17] P. Khuc Trong, H. Doerflinger, J. Dunkel, D. St. Johnston, and R.E. Goldstein, Cortical Microtubule Nucleation Can Organise the Cytoskeleton of *Drosophila* Oocytes to Define the Anteroposterior Axis, eLife 4, e06088 (2015).

[18] W. Lu, M. Lakonishok, A.S. Serpenskaya, D. Kirchenbuechler, S.-C. Ling and V.I. Gelfand, Ooplasmic flow cooperates with transport and anchorage in *Drosophila* oocyte posterior determination, J. Cell. Biol. 217, 3497 (2018).

[19] F. Gittes, B. Mickey, J. Nettleton and J. Howard, Flexural Rigidity of Microtubules and Actin Filaments Measured from Thermal Fluctuations in Shape, J. Cell Biol. 120, 923 (1993).

[20] K. Visscher, M.J. Schnitzer, and S.M. Block, Single Kinesin Molecules Studied With a Molecular Force Clamp, Nature 400, 184 (1999).

[21] G. Herrmann and R.W. Bungay RW, On the stability of elastic systems subjected to nonconservative forces, J. Appl. Mech. 31, 435 (1964).

[22] P.V. Bayly and S.K. Dutcher, Steady dynein forces induce flutter instability and propagating waves in mathematical models of flagella, J. R. Soc. Interface 13, 20160523 (2016).

[23] G. De Canio, E. Lauga, and R.E. Goldstein, Spontaneous oscillations of elastic filaments induced by molecular motors, J. R. Soc. Interface 14, 20170491 (2017).

[24] F. Ling, H. Guo, and E. Kanso, Instability-driven oscillations of elastic microfilaments, J. R. Soc. Interface 15, 149 (2018).

[25] Y.-N. Young and M.J. Shelley, Stretch-Coil Transition and Transport of Fibers in Cellular Flows, Phys. Rev. Lett. 99, 058303 (2007).

[26] V. Kantsler and R.E. Goldstein, Flucutations, Dynamics, and the Stretch-Coil Transition of Single Actin Filaments in Extensional Flows, Phys. Rev. Lett. 108, 038103 (2012).

[27] D.B. Stein and M.J. Shelley, Coarse graining the dynamics of immersed and driven fiber assemblies, Phys. Rev. Fluids 4, 073302 (2019).

[28] I.M. Palacios and M. Drechsler, private communication (2020), based on methods detailed earlier [11, 16].

[29] W. Lu, M. Winding, M. Lakonishok, J. Wildonger and V.I. Gelfand, Microtuble-microtubule sliding by kinesin-1 is essential for normal cytoplasmic streaming in *Drosophila* oocytes, Proc. Natl. Acad. Sci. USA 113, E4995 (2016).

[30] J.B. Keller and S.I. Rubinow, Slender-body theory for slow viscous flow, J. Fluid Mech. 75, 705 (1976).

[31] J. Gray and G.J. Hancock, The propulsion of sea-urchin spermatozoa, J. Exp. Biol. 32, 802 (1955).

[32] A.K. Tornberg, and M.J. Shelley, Simulating the dynamics and interactions of flexible fibers in Stokes flows, J. Comput. Phys. 196, 1 (2004).

[33] R.E. Goldstein and S.A. Langer, Nonlinear dynamics of stiff polymers, Phys. Rev. Lett. 75, 1094 (1995).

[34] See Supplemental Material at http://link.aps.org/supplemental/xxx for further details and results.

[35] Further examples and details are in: G. De Canio, Motion of filaments induced by molecular motors: from individual to collective dynamics, PhD thesis, University of Cambridge (2018).

[36] C. Maul and S. Kim, Image systems for a Stokeslet inside a rigid spherical container, Phys. Fluids 6, 2221 (1994).

[37] R.E. Goldstein, Green Algae as Model Organisms for Biological Fluid Dynamics, Annu. Rev. Fluid Mech. 47, 343 (2015).

[38] T. Niedermayer, B. Eckhardt and P. Lenz, Synchronization, phase locking, and metachronal wave formation in ciliary chains, Chaos 18, 037128 (2008).

[39] Y. Yotsuyanagi, Recherches sur les phénomenès moteurs dans les fragments de protoplasme isolés. I. Mouvement rotatoire et le processus de son apparition, Cytologia 18, 146 (1953)

40. Recherches sur les phénomenès moteurs dans les fragments de protoplasme isolées. II. Mouvements divers d’etermin’es par la condition de milieu, Cytologia 18, 202 (1953).

[40] H. Wioland, F.G. Woodhouse, J. Dunkel, J.O. Kessler, and R.E. Goldstein, Confinement Stabilizes a Bacterial Suspensions into a Spiral Vortex, Phys. Rev. Lett. 110, 268102 (2013).

[41] D. Saintillan and M.J. Shelley, Instabilities and Pattern Formation in Active Particle Suspensions: Kinetic Theory and Continuum Simulations, Phys. Rev. Lett. 100, 178103 (2008).

[42] D. Saintillan, M. J. Shelley, A. Zidovska, Extensile motor activity drives coherent motions in a model of interphase chromatin, Proc. Natl. Acad. Sci. USA 115, 11442 (2018).

[43] F.G. Woodhouse and R.E. Goldstein, Spontaneous Circulation of Confined Active Suspensions, Phys. Rev. Lett. 109, 168105 (2012).

[44] E. Nazockdast, A. Rahimian, D. Zorin, and M.J. Shelley, A fast platform for simulating semi-flexible fiber suspensions applied to cell mechanics, J. Comput. Phys. 329 (2017).

[45] Differentiation of (2) yields an equivalent equation for the tangent-vector field **r**_s_ [27], which we have found to be numerically more stable.

[46] L.D. Landau and E.M. Lifshitz, Theory of Elasticity, 2nd ed. (Pergamon Press, Oxford, 1970), p. 99, Problem 7.

[47] R.E. Goldstein, P.B. Warren and R.C. Ball, Shape of a Ponytail and the Statistical Physics of Hair Fiber Bundles, Phys. Rev. Lett. 108, 078101 (2012).

[48] P. Loiseau,, R. Davies, L.S. Williams, M. Mishima and I.M. Palacios, *Drosophila* PAT1 is required for Kinesin-1 to transport cargo and to maximize its motility, Development 137, 2763 (2010).

[49] D.R. Brumley, K.Y. Wan, M. Polin and R.E. Goldstein, Flagellar synchronization through direct hydrodynamic interactions, eLife 3, e02750 (2014).

[50] J.R. Blake, A note on the image system for a stokeslet in a no-slip boundary, Math. Proc. Camb. Phil. Soc. 70, 303 (1971).

[51] J.-B. Thomazo, E. Lauga, B. Le Révérend, E. Wandersman, and A.M. Prevost, Collective stiffening of soft hair assembles, arXiv:2002.02834.

