## Supplemental Material for "Swirling Instability of the Microtubule Cytoskeleton"

(Dated: August 27, 2020)

This file contains calculational details for computations of discrete filaments and the continuum model, further results from linear stability analyses in the latter, and quantitative analysis of streaming speeds in the swirling state.

### I. CALCULATIONAL DETAILS FOR DISCRETE FILAMENTS

#### A. Image system inside a sphere

By modelling the motor-cargo ensemble with fluid-entraining follower forces and approximating the system geometry by a spherical container, we take advantage of the analytical solution [S1] for the velocity field of a point force inside a sphere with a no-slip wall. These simplifications allow us to compute the background flow directly, instead of numerically solving the Stokes equations for the fluid flow. For a point force of magnitude $F$ located at $\mathbf{x}_0$ inside a sphere of unit radius, the $\mathrm{m}^{\mathrm{th}}$ component of the velocity is

$$u_{\mathrm{m}} = \frac{F_{\mathrm{j}}}{8\pi\mu}\left[G_{\mathrm{jm}}\left(\mathbf{x}-\mathbf{x}_0\right) + \bar{G}_{\mathrm{jm}}\left(\mathbf{x}\right)\right] , \quad \text{(S1)}$$

where $\mu$ is the dynamic viscosity of the fluid, $G_{\mathrm{jm}}$ is the Green's function

$$G_{\mathrm{jm}}\left(\mathbf{x}-\mathbf{x}_0\right) = \frac{\delta_{\mathrm{jm}}}{r} + \frac{\left(x_{\mathrm{j}}-x_{0_{\mathrm{j}}}\right)\left(x_{\mathrm{m}}-x_{0_{\mathrm{m}}}\right)}{r^3} , \quad \text{(S2)}$$

and

$$\begin{aligned}\bar{G}_{\mathrm{jm}}\left(\mathbf{x}\right) = {} & \hat{e}_{\mathrm{m}}\hat{e}_{\mathrm{k}}\left\{\frac{1-3R_0^2}{2R_0^3}\,G_{\mathrm{jk}}\left(\mathbf{x}-\bar{\mathbf{x}}_0\right) - \frac{1-R_0^2}{R_0^4}\,\hat{e}_{\mathrm{l}}\,G_{\mathrm{jk},l}\left(\mathbf{x}-\mathbf{x}_0\right) - \frac{(1-R_0^2)^2}{4R_0^5}\,\nabla^2 G_{\mathrm{jk}}\left(\mathbf{x}-\bar{\mathbf{x}}_0\right)\right\} \\ & + \left(\delta_{\mathrm{km}}-\hat{e}_{\mathrm{k}}\hat{e}_{\mathrm{m}}\right)\left\{\frac{3R_0^2-5}{2R_0^3}\,G_{\mathrm{jk}}\left(\mathbf{x}-\bar{\mathbf{x}}_0\right) + \frac{(1-R_0^2)^2}{4R_0^5}\,\nabla^2 G_{\mathrm{jk}}\left(\mathbf{x}-\bar{\mathbf{x}}_0\right)\right\} \\ & + \frac{1-R_0^2}{R_0^4}\,\hat{e}_{\mathrm{k}}\left(\delta_{\mathrm{lm}}-\hat{e}_{\mathrm{l}}\hat{e}_{\mathrm{m}}\right)G_{\mathrm{jk},l}\left(\mathbf{x}-\bar{\mathbf{x}}_0\right) - \frac{3(R_0^2-1)}{R_0^3}\frac{\left(\delta_{\mathrm{jm}}-\hat{e}_{\mathrm{j}}\hat{e}_{\mathrm{m}}\right)}{\bar{r}} + \left(R^2-1\right)\left(\delta_{\mathrm{km}}-\hat{e}_{\mathrm{k}}\hat{e}_{\mathrm{m}}\right)\frac{\partial\varphi_{\mathrm{k}}}{\partial x_{\mathrm{j}}} , \end{aligned} \quad \text{(S3)}$$

with

$$\varphi_{\mathrm{k}} = -\frac{3(R_0^2-1)}{2R_0^3}\,\frac{x_{\mathrm{k}}}{\bar{r}}\,\frac{R-\bar{R}_0\cos\alpha+\bar{r}\cos\alpha}{R\bar{R}_0^2\sin^2\alpha} . \quad \text{(S4)}$$

Here, $R_0$ and $\bar{R}_0$ are the norms of $\mathbf{x}_0$ and its image $\bar{\mathbf{x}}_0$, respectively, $\hat{\mathbf{a}} = \mathbf{x}_0/R_0$ is the unit vector in the axial direction, $r = |\mathbf{x}-\mathbf{x}_0|$, $\bar{r} = |\mathbf{x}-\bar{\mathbf{x}}_0|$, $R$ is the norm of $\mathbf{x}$, and $\alpha$ is the angle between $\mathbf{x}$ and $\hat{\mathbf{a}}$.

#### B. Geometrical formulation

For both the discrete-filament computations and the self-consistent model, we use the tangent angle representation of the filament. For a filament of length $L$, parameterized by arclength $s$, anchored at wall in the $x-y$ plane, with $z$ orthogonal to the plane, into the fluid, the position vector is $\mathbf{r}(s) = (x(s), z(s))$, and the unit tangent and normal vectors $\hat{\mathbf{t}}(s) = \mathbf{r}_s$ and $\hat{\mathbf{n}}(s)$ are

$$\hat{\mathbf{t}}(s) = \cos\theta(s)\hat{\mathbf{e}}_x + \sin\theta(s)\hat{\mathbf{e}}_z \quad \text{(S5)}$$

$$\hat{\mathbf{n}}(s) = \sin\theta(s)\hat{\mathbf{e}}_x - \cos\theta(s)\hat{\mathbf{e}}_z , \quad \text{(S6)}$$

where $\theta(s)$ is the angle the tangent vector makes with respect to the $x$-axis. The curve is obtained by integration of $\hat{\mathbf{t}}$, and assuming $(x(0), z(0)) = (0,0)$, we have

$$x(s) = \int_0^s ds'\cos\theta(s') , \quad z(s) = \int_0^s ds'\sin\theta(s') . \quad \text{(S7)}$$

It follows from the above that

$$\frac{\partial \hat{\mathbf{t}}}{\partial t} = -\theta_t \hat{\mathbf{n}} \ , \tag{S8}$$

from which we can obtain the equation of motion for $\theta$ given that for $\mathbf{r}$.

The equation of motion is that of elastohydrodynamics within resistive force theory,

$$\left(\zeta_{\|}\hat{\mathbf{t}}\hat{\mathbf{t}} + \zeta_{\perp}\hat{\mathbf{n}}\hat{\mathbf{n}}\right)\cdot(\mathbf{r}_t - \mathbf{U}) = -A\mathbf{r}_{ssss} - (\Lambda \mathbf{r}_s)_s \ , \tag{S9}$$

where $A$ is the filament bending modulus, $\Lambda$ is the Lagrange multiplier, and $\mathbf{U} = U\hat{\mathbf{e}}_x + V\hat{\mathbf{e}}_z$ is the background flow experienced by the filament. We express the derivatives in (S9) in terms of the curvature $\kappa = \theta_s$, using the Frenet-Serret equations $\hat{\mathbf{t}}_s = -\kappa\hat{\mathbf{n}}$ and $\hat{\mathbf{n}}_s = \kappa\hat{\mathbf{t}}$, and rescale the equation of motion via

$$\mathbf{r} = L\mathbf{r}' \ , \quad z = Lz' \ , \quad s = Ls' \ , \quad \kappa = \frac{1}{L}\kappa' \ ,$$
$$\Lambda = \frac{A}{L^2}\Lambda' \ , \quad t = \frac{\zeta_\perp L^4}{A}t' \ , \quad U = \frac{\zeta_\perp L^3}{A}U'. \tag{S10}$$

If we let $\zeta_\perp/\zeta_\| \simeq 2$, and use $\hat{\mathbf{e}}_x = \cos\theta\hat{\mathbf{t}} + \sin\theta\hat{\mathbf{n}}$ and $\hat{\mathbf{e}}_z = \sin\theta\hat{\mathbf{t}} - \cos\theta\hat{\mathbf{n}}$, and then drop the primes, we obtain

$$\mathbf{r}_t = (U\cos\theta + V\sin\theta)\,\hat{\mathbf{t}} + (U\sin\theta - V\cos\theta)\,\hat{\mathbf{n}} + \left(\kappa_{ss} - \kappa^3 + \kappa\Lambda\right)\hat{\mathbf{n}} + \eta\left(3\kappa\kappa_s - \Lambda_s\right)\hat{\mathbf{t}} \ . \tag{S11}$$

Differentiating (S11) with respect to $s$ to obtain the equation of motion of the tangent angle $\theta$ and that for $\Lambda$:

$$\theta_t = -\theta_{ssss} + 3\theta_s^2\theta_{ss} - (\theta_s\Lambda)_s + 2\left(3\theta_s^2\theta_{ss} - \theta_s\Lambda_s\right) - U_s\sin\theta + V_s\cos\theta \ , \tag{S12}$$

while enforcing inextensibility leads to the equation for $\Lambda$,

$$\left(\partial_{ss} - \frac{1}{2}\theta_s^2\right)\Lambda = -\frac{1}{2}\theta_s^4 + 3\theta_{ss}^2 + \frac{7}{2}\theta_s\theta_{sss} + \frac{M}{\eta}\sin(\theta)\cos(\theta) \ . \tag{S13}$$

The boundary conditions at the attachment point are those of a clamped filament,

$$\theta(0) = \frac{\pi}{2} \ , \quad \theta_{sss}(0) - \theta_s(0)^3 + \theta_s(0)\Lambda(0) = 0 \ , \tag{S14}$$
$$\Lambda_s(0) - 3\theta_s(0)\theta_{ss}(0) = 0 \ .$$

whereas at the free end we have

$$\theta_s(1) = 0 \ , \quad \theta_{ss}(1) = 0 \ , \quad \Lambda(1) = 0 \ . \tag{S15}$$

#### C. Self-consistent model

If we take the typical bent filament shape in the streaming regime (Fig. 2(d) of main text) and place it near a flat no-slip wall, we can understand in the simplest situation the flow it produces. As shown in the streamlines and velocity colormap of Figure S1 there is a boundary-layer phenomenon near the wall that arises from the combination of tangential forcing of the flow by the motors and the no-slip condition at the wall. The most prominent part of the flow is a lobe of high speeds emanating from the bent portion of the filament, directed downstream. This feature forms the basis of the self-consistent model for the swirling transition.

As Fig. S1 shows, the flow downstream from a bent filament is approximately simple shear. This can be seen directly in Blake's analysis [S2] of the flow due to a point force near a no-slip wall. For the purposes of a simple self-consistent model, we consider only the asymptotic form of that flow evaluated at $\mathbf{x} = (x, y, z)$ due to a point force $\mathbf{F}$ at $(0, 0, h)$:

$$u_i \simeq \frac{F_k}{8\pi\mu}\left[\frac{12hx_ix_\alpha x_3\delta_{k\alpha}}{|\mathbf{x}|^5}\right. \tag{S16}$$
$$\left. + h^2\delta_{k3}\left(-\frac{(12+6\delta_{i3})x_ix_3}{|\mathbf{x}|^5} + \frac{30x_ix_3^3}{|\mathbf{x}|^7}\right)\right] .$$

Taking the leading-order term, and $\mathbf{F}$ along $x$, we obtain

$$\mathbf{u}(x,z) = \dot{\gamma}(x)z\hat{\mathbf{e}}_x \tag{S17}$$

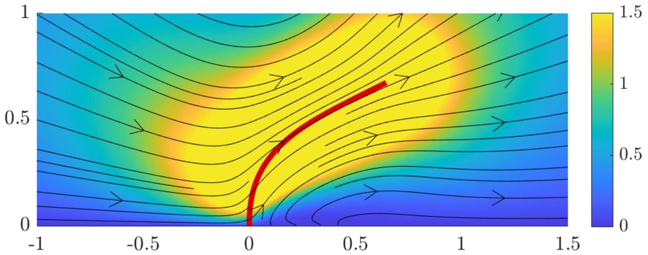


FIG. S1: Flow field near a filament in the swirling regime. A filament in the bent shape found in the spherical geometry of the main text was placed near a flat, no-slip wall and the resulting flow field computed numerically. Note the shear flow downstream.

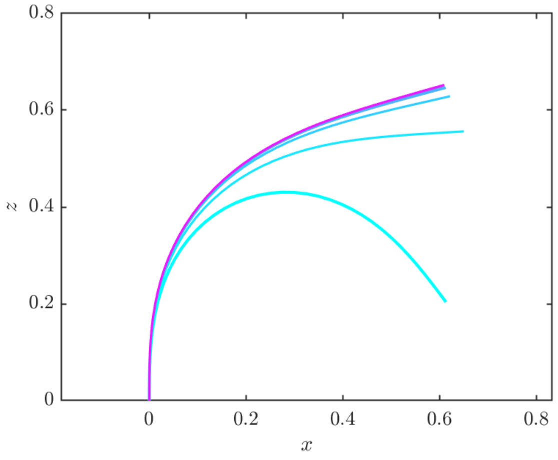


FIG. S2: Results of the self-consistent calculation. Relaxation of a filament to steady bent configuration, at $\sigma = 45$ and $\rho = 5$, with colors interpolating between highly bent initial condition (cyan) to final state (magenta) over a (dimensionless) time of 0.05.

where the shear rate is

$$\dot{\gamma}(x) = \frac{3Fh}{2\pi\mu x^3} \,. \tag{S18}$$

In the notation used in the tangent angle dynamics (S12), we have $U = Mz(s)$ and $V = 0$, where

$$M = \frac{\eta\dot{\gamma}L^4}{A} \sim \frac{3\sigma}{c}\left(\frac{\rho}{\phi}\right)^{3/2} \,. \tag{S19}$$

a control parameter that appears also in the dynamics elastic filaments in extensional flows [S3, S4]. Here, the second relation uses the definitions of the dimensionless control parameters $\sigma$ and $\rho$ from the main text. To complete the self-consistency calculation, we set the $x$-component of the force $\mathbf{F}$ to be proportional to the projection of the tangent vector at the distal tip, so $M \to M\sin(\theta(L))$

The tangent angle dynamics (S12) and (S13) were solved by a Crank-Nicholson method via a pentadiagonal matrix for the fourth derivative, treating all the nonlinearities explicitly, and discretizing the $\Lambda$ problem as a tridiagonal matrix in which the diagonal term involving $\theta_s^2$ was updated after each time step. The initial condition was $\theta(0,s) = \pi/2 - a\sin(\pi s/2)^2$, with $a = 2.5$ and the time step $dt = 0.1ds^4$, with $ds = 1/(N-1)$, where $N$ is the total number of grid points ($N = 41$ is sufficient).

Figure S2 shows a typical result of the self-consistent calculation in the regime of stable streaming, illustrating how a highly bent initial condition relaxes to a conformation like that seen in the calculations described in the main text (compare Figs. 3(c) from discrete filament calculations and 4(b) from continuum model).

### II. FURTHER RESULTS FROM CONTINUUM MODEL

#### A. Dynamics of the transition to swirling

In Supplemental Video 1, we show the dynamics of the transition from the unstable equilibrium of radially oriented filaments to stable streaming flow, corresponding to Fig. 4b (main text). In this parameter regime, the filament array initially buckles, and large azimuthal flows are generated by rapid deformation of the array. As the filament motion slows, the filaments become oriented parallel to the cylinder walls. Azimuthal flows are continuously generated by the streaming mechanism, and the drag from these flows moving past the stationary filaments is sufficient to pin them into the observed conformation.

#### B. Effect of cylindrical confinement

In the main text, we present the evolution equation for a small perturbation to the equilibrium solution in a planar geometry. This equation:

$$g_t = -g_{zzzz} - \sigma\left[(1-z)g_z\right]_z + \rho\left[\sigma(1-z)g + g_{zz}\right], \tag{S20}$$

remains local due to the simplicity of the Stokes flow, allowing the individual terms to be easily identified and interpreted. In order to better capture the real geometry of the oocyte, in the main text we present the results of the linear stability analysis in a cylindrical geometry (Fig 4a, main text). The same analysis, over the same range of effective densities ($\rho$) and effective motor force-densities ($\sigma$) is shown in Fig. S3, computed for the planar geometry. These results are qualitatively the same as those shown in the main text in Fig 4a, indicating that the essential structure of the swirling transition is due to the interactions between nearby filaments in the fiber

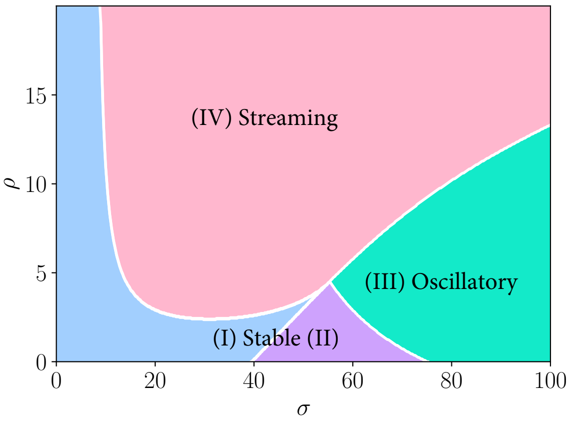


FIG. S3: Results of the stability analysis in the planar geometry. Regions are colored and coded as in the main text.

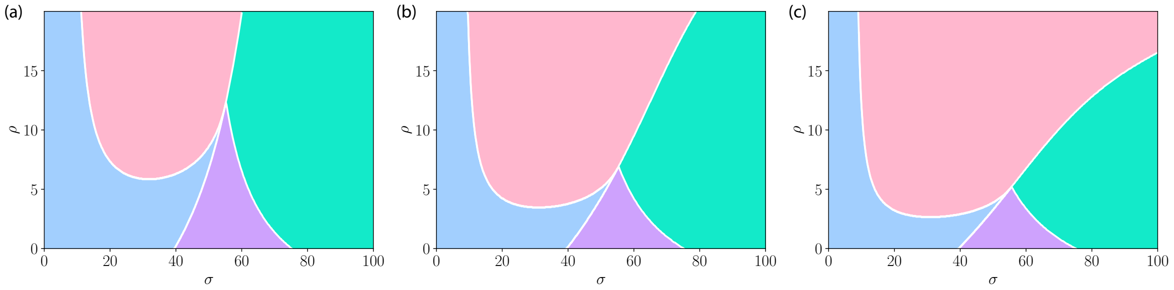


FIG. S4: Results of the stability analysis in the confined geometry. The confinement ratios are 2.2:1 (Panel a), 4:1 (Panel b), and 20:1 (Panel c). See Fig. S3 for a description of the different regions.

bed, and does not depend in a critical way on the specific geometry of the oocyte.

In, Fig. S4 we present the same analysis for different confinement ratios — 2.2:1 (Fig S4a), 4:1 (Fig S4b), and 20:1, (Fig S4c). Note that the ratio 2.2:1 indicates a cylinder diameter of $2.2L$, which is extremely confined, with filaments reaching $\approx$ 91% of the way to the center of the cylinder. Even under such strong confinement, the qualitative structure remains, although the quantitative value at which the bifurcations occur changes significantly. The effect of strong confinement is to suppress the appearance of streaming at larger forcing values. This makes sense: when forcing is large, the tendency of lone fibers is to oscillate. In the planar system, when fibers are sufficiently dense, the strong flows caused by motors walking on neighboring fibers is sufficient to pin the fiber into a stable deformed state, suppressing oscillations (in fact, the drag from these flows produces extensile forces on the fibers, reducing the net contractile forcing generated by the motors). Under high confinement, the strength of such flows near the fiber tips is reduced, and the flow can no longer suppress the tendency of the fiber to oscillate. When the confinement is not so extreme, even the quantitative values change little, from 10:1 (Fig 4a, main text), to 20:1 (Fig S4c), to the flat geometry (Fig S3).

It may surprise the reader that when $\rho \to 0$, the behavior (as a function of $\sigma$) at all confinement ratios converges to *exactly* the same thing. This is because the continuum model imposes boundary conditions on the coarse-grained flow only; as $\rho \to 0$ this flow goes to 0 with it and the dynamics are those of a fiber moving in a quiescent background.

#### C. Streaming Speeds in Dense Arrays

By solving for $-\mathbf{r}_{ssss} + (\Lambda \mathbf{r}_s)_s$ in Eq. 2 (main text), substituting into Eq. 3 (main text), defining $\xi \equiv \chi_{\mathrm{MT}} \rho \mathcal{J}^{-1}$, and ignoring frame transformations for the sake of simplicity, we find that:

$$-\nabla^2 \mathbf{u} + \nabla p + \xi(\mathbb{I} - \mathbf{r}_s\mathbf{r}_s/2)(\mathbf{u} - \mathbf{r}_t) = \xi\sigma\mathbf{r}_s. \qquad \text{(S21)}$$

This is a forced Brinkman equation with an anisotropic permeability. When $\xi$ is large, the skeletal drag term dominates the left-hand side, and at steady state $\mathbf{r}_t = 0$. We may thus approximate the steady streaming velocity as simply $\mathbf{u} \approx \sigma(\mathbb{I} + \mathbf{r}_s\mathbf{r}_s)\mathbf{r}_s = 2\sigma\mathbf{r}_s$. For homogeneous flows in the planar geometry, and axisymmetric flows in the cylindrical geometry, the flow is purely in the $\hat{x}$ or $\hat{\theta}$ (azimuthal) direction, respectively. Since $\mathbf{r}_s$ is not purely coincident with $\hat{x}$ or $\hat{\theta}$, some of the forcing generated by the molecular motors must be absorbed into the pressure gradient, and so we take this estimate to be an upper bound: $v_{\mathrm{streaming}} \leq 2\sigma$. The maximum steady-state streaming speed as a function of $\sigma$, as computed by the continuum model in 10:1 confinement, is shown in Fig. S5, for $\rho = 10, 20, \ldots, 50$. For smaller values of $\rho$, the fiber is less deformed and drag contributes less to the balance in Eq. S21, leading to a relationship closer to $v_{\mathrm{streaming}} = \sigma$. When $\rho$ is larger, drag dominates Eq. S21, the fiber is nearly azimuthally aligned, and the streaming speed approaches the bound $v_{\mathrm{streaming}} = 2\sigma$.

In dimensional units, this bound is $v_{\mathrm{streaming}} \leq 2\sigma A/\eta L^3$. In the main text, we used a simple argument based on Darcy's law to estimate $v_{\mathrm{streaming}} =$

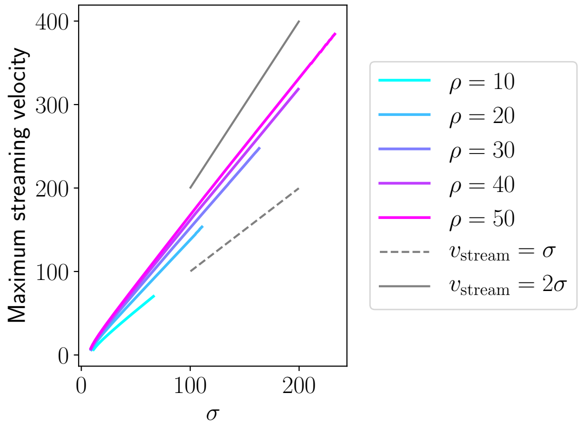


FIG. S5: Dependence of streaming speed on the force density ($\sigma$) generated by kinesin-1 motors. When $\rho$ is large, the streaming speed approaches the estimate $v_{\mathrm{streaming}} = 2\sigma$.

$(\sigma A/\eta L^3)(8\pi/c)$. The leading factor of two in the more refined estimate presented in this SI comes from taking into account fiber anisotropy and the nearly azimuthal alignment of the fibers, while the disappearance of the factor of $8\pi/c$ arises from properly accounting for the fibers aspect ratio. In either case, the disappearance of the MT density from the estimate of streaming velocity arises due to the fact that both the forcing $(\xi\sigma\mathbf{r}_s)$ and the skeletal drag $(\xi(\mathbb{I}-\mathbf{r}_s\mathbf{r}_s/2))$ scale with the density $\rho$.


[S1] C. Maul and S. Kim, Image systems for a Stokeslet inside a rigid spherical container, Phys. Fluids **6**, 2221 (1994).

[S2] J.R. Blake, A note on the image system for a stokeslet in a no-slip boundary, Math. Proc. Camb. Phil. Soc. **70**, 303 (1971).

[S3] Y.-N. Young and M.J. Shelley, Stretch-Coil Transition and Transport of Fibers in Cellular Flows, Phys. Rev. Lett. **99**, 058303 (2007).

[S4] V. Kantsler and R.E. Goldstein, Flucutations, Dynamics, and the Stretch-Coil Transition of Single Actin Filaments in Extensional Flows, Phys. Rev. Lett. **108**, 038103 (2012).
